# Temporal stability of human sperm mosaic mutations results in life-long threat of transmission to offspring

**DOI:** 10.1101/2020.10.14.339796

**Authors:** Xiaoxu Yang, Martin W. Breuss, Xin Xu, Danny Antaki, Kiely N. James, Valentina Stanley, Laurel L. Ball, Renee D. George, Sara A. Wirth, Beibei Cao, An Nguyen, Jennifer McEvoy-Venneri, Guoliang Chai, Shareef Nahas, Lucitia Van Der Kraan, Yan Ding, Jonathan Sebat, Joseph G. Gleeson

## Abstract

Every newborn harbors scores of new single nucleotide variants (SNVs) that may impact health and disease^1–4^; the majority of these are contributed by the paternal germ cells^5^. In some cases, these mutations are identifiable in a subset of the parents’ cells—a phenomenon called mosaicism, which is capable of producing disease recurrence^6–8^. Here, we provide a comprehensive analysis of male gonadal mosaic mutations, employing 300× whole genome sequencing (WGS) of blood and sperm in 17 healthy individuals, including assessment across multiple samples and age groups. Approximately 1 in 15 healthy males is predicted to harbor a transmissible, likely pathogenic exonic variant that is mosaic in his sperm. In general, only a third of sperm mosaic mutations were detectable in blood cells, all were remarkably stable over the course of months to years, and 23% were present in 5% or more of sperm cells. There was no evidence of age-dependent clonal expansion or collapse, as seen in hematopoiesis. Thus, despite the observed increase of mutations in offspring of men with advanced paternal age, detectable sperm mosaicism remains stable, represents a life-long transmission risk to offspring, and suggests a testis stem cell niche that prevents widespread clonality.

## Main

Cellular proliferation and metabolism introduce mutations into the genome of every cell^9–12^. If these occur in embryogenesis, they may spread widely across or within tissues at appreciable allelic fractions (AF; i.e. fraction of mutant DNA molecules). These mutations are unable to transmit to offspring unless they are present within primordial germ cells (PGCs)^9,13,14^. We previously distinguished three types of sperm mosaicism^7^: ‘Type I’, occurring during or after the terminal spermatogonial stem cell division, with no chance of recurrence; ‘Type II’, occurring in spermatogonial stem cells, accumulating with paternal age, and possibly undergoing positive selection (i.e. ‘selfish sperm’ model)^15^; and ‘Type III’, occurring during embryogenesis of the father, contributing to PGCs and potentially other tissues. We found that Type III accounts for ~4% of paternally-phased mutations detected in offspring, in agreement with previous indirect estimates^6,14,16^.

Here, we study the landscape of sperm mosaicism to determine the number of transmissible mosaic SNVs (mSNVs) in healthy men, temporal stability within an individual, and changes in old vs. young men. We used >300× WGS from blood and sperm in 12 males aged 18-22 years (young age; YA, ID01-12) to establish baseline mosaicism before any clonal selection might change abundance (Fig. 1a, Extended Data Fig. 1). We further collected multiple (up to 3) sperm samples for 9 of these individuals at ~6 month intervals. Finally, we assessed a cohort of 5 males aged 48-62 years (advanced age; AA, ID13-17). Together, these approaches revealed the inter-subject, intra-subject, and age-dependent variability of sperm and blood mosaicism.

**Figure 1.**
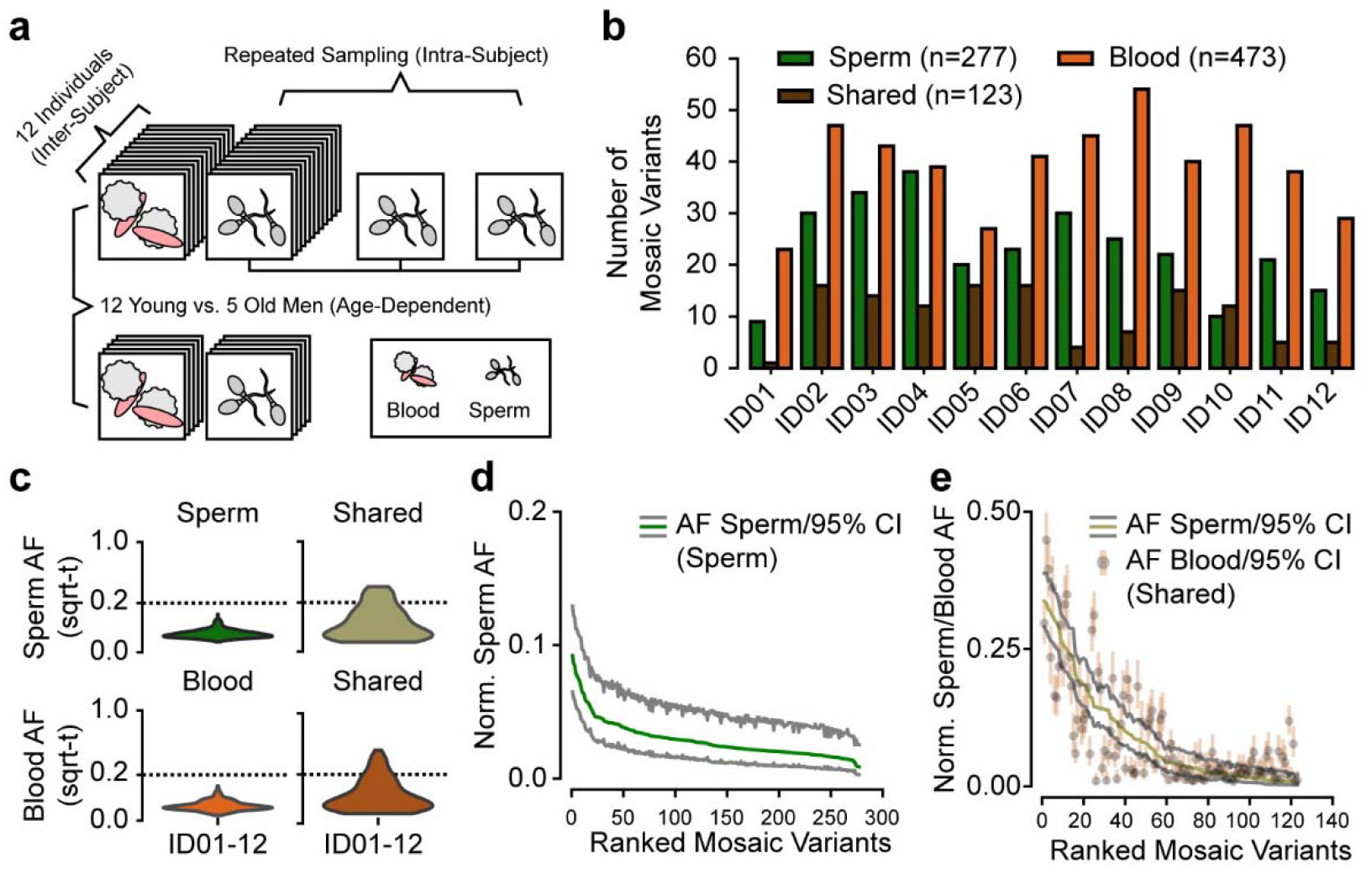
Analysis in 12 young aged men uncovers the landscape of sperm mosaicism. **a**, Sampling strategy: 12 healthy males of young age (YA, 18-22 years, blood and up to 3 sperm samples) and 5 healthy males of advanced age (AA, >48 years, blood and 1 sperm sample). Samples subjected to 300× whole-genome sequencing (WGS). **b**, Number of detected mosaic variants per individual from each class (*Sperm, Shared, Blood*), typically there are more *Blood* than *Shared* or *Sperm*. **c**, AF distribution (square root-transformed; sqrt-t) of *Sperm, Shared*, and *Blood* variants in the entire YA cohort. *Shared* variants show higher peak and overall AF compared to *Sperm* and *Blood*. **d-e**, Ranked plot of the estimated sperm and blood AF with 95% confidence intervals (exact binomial CIs) from the YA cohort, grouped by the classes described above. *Sperm* (d) variants showed steeper decay curves, indicating a relative lower mutation or higher expansion rate. *Shared* variants (e) showed a shallower decay and higher AF, indicating a different speed for the accumulation of mutations during early embryonic development.

### Detectable mosaicism is more common within than across tissues

We employed a combination of state-of-the-art mosaic variant callers for mSNVs and mINDELs (small insertions and deletions), termed 300×MSMF (Extended Data Fig. 2a)^7,17,18^. This approach, which identifies both tissue-specific and tissue-shared variants, demonstrated sensitivity to ~1% AF and a validation rate of 97.5% on benchmarked data (Extended Data Fig. 2b-d).

We found that each YA male harbored between 9-38 sperm-specific (‘*Sperm*’) (mean ± SD: 23.1 ± 9.0; total: 277), 1-16 tissue-shared (‘*Shared*’) (10.3 ± 5.5; 123), and 23-54 blood-specific (‘*Blood*’) (39.4 ± 9.1; 473) variants (Fig. 1b, Extended Data Fig. 3a-c, Supplementary Dataset 1). Thus, an average of >30 variants were detected in sperm with an average AF of 4.8%; two-thirds of these were found exclusively in semen. Conversely, 80% of detected mosaic variants from blood were not identified in sperm, and thus likely not transmissible. The AFs of the *Shared* variants were higher than tissue-specific ones (Fig. 1c-e, Extended Data Fig. 3d-e) and correlated tightly in sperm and blood (Extended Data Fig. 3f). These results suggest an early developmental origin of *Shared* variants and a continuous accumulation of sperm- and blood-specific variants after progenitor lineage separation.

### Modest sperm mosaicism variation within an individual

For 9 YA individuals, we obtained samples to measure the stability of sperm mosaicism over the course of one year (Fig. 2a). First, to assess whether new mosaic variants appeared over time, we performed 300×MSMF on two additional semen samples from ID04 and ID12 (Fig. 2b). Second, to accurately quantify the stability of mosaic variants, we performed targeted amplicon sequencing (TAS) for ~50% of *Sperm* and *Shared* candidate variants in sperm samples from YA individuals from up to two additional time points (Fig. 2c).

**Figure 2.**
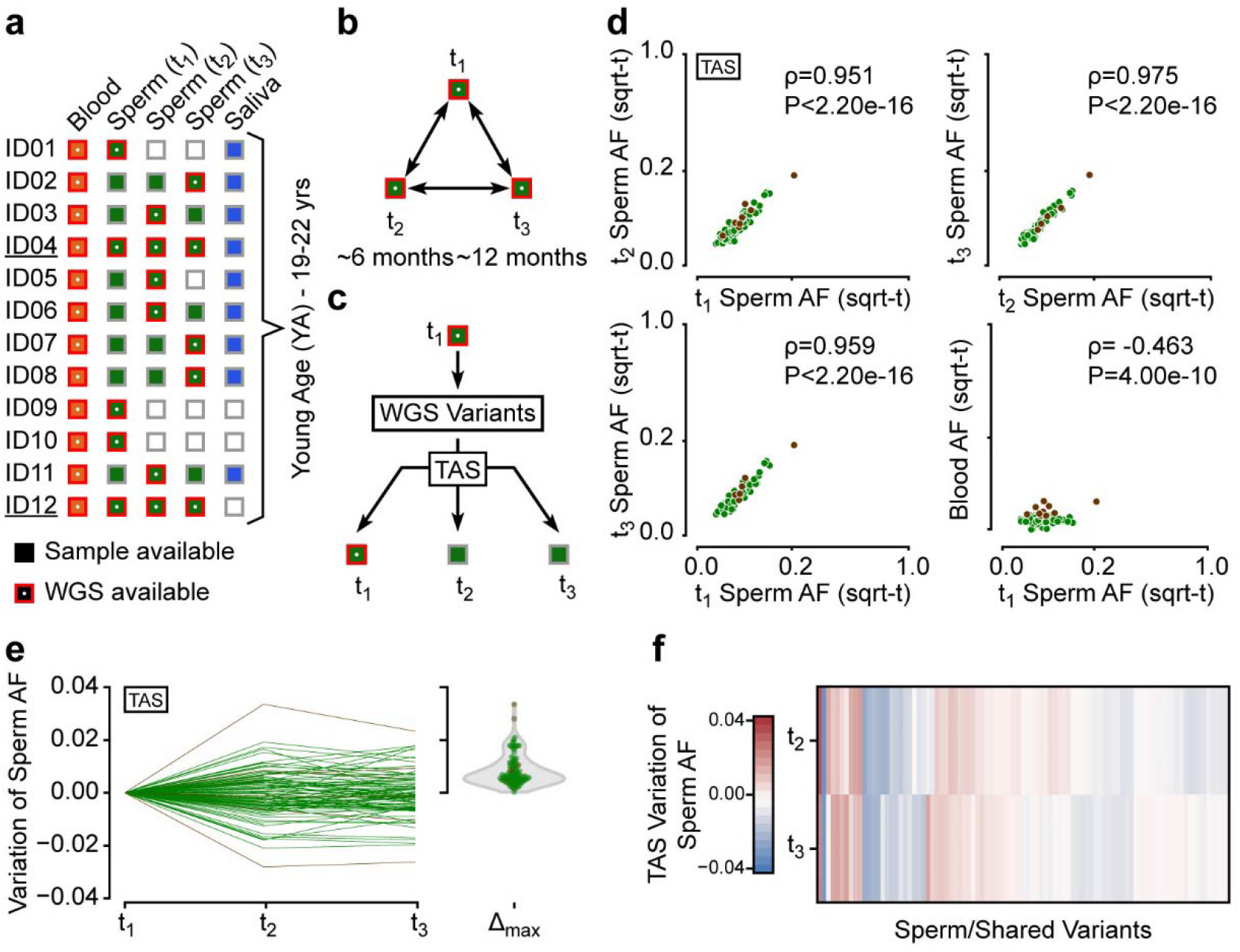
Sperm mosaic variants showed temporal stability within an individual. **a**, Available blood, sperm, and saliva samples for ID01-12 and their WGS status. ID04 and ID12 had three samples subjected to WGS. **b**, Analysis strategy for ID04 and ID12. **c**, Assessment of a subset of sperm mosaic variants called by WGS through targeted amplicon sequencing (TAS) in all available samples of an individual. Note that TAS typically has >5000×, resulting in increased sensitivity. **d,** Pair-wise AF comparison of sperm mosaic variants across the YA cohort by TAS. All tested variants were detected in all available sperm samples. Each plot shows Spearman’s ρ and P-value across all variants. **e**, Sperm AF changes for each tested variant. Variation was typically below 0.02.Violin plot shows the maximal, absolute change for each variant. **f**, Heatmap of AF variation relative to t_1_ for variants with 3 available samples. No patterns of clear linear increase or decrease were observed.

In the first approach, we detected 125 variants from ID04’s and ID12’s respective three semen samples. The AF of variants that were mosaic in sperm correlated tightly across time points (Extended Data Fig. 4a). For a number of variants that were close to the detection limit of WGS, we observed several that were absent in one or two of the sperm datasets, but in general new somatic variants did not appear or drop out.

Second, we employed TAS on sperm, blood, and saliva samples, with average readdepths >5000×. We confirmed 130 variants as mosaic in sperm, blood, or both, and subsets of these had two (n=105) or three (n=91) sperm or saliva (n=41) samples available (Supplementary Dataset 2). Saliva was tightly correlated to blood, consistent with leukocytes being a main sources of salivary DNA^19^ (Extended Data Fig. 4b).

All mutations observed in sperm through TAS were detectable and correlated tightly across all semen samples within a subject (Fig. 2d). Variants were stable across one year, as AF change was typically below 2% (Fig. 2e). The variation across time points, however, was imperfectly correlated with the initially observed AF, and fold-change was as high as 1.5-2 for low AF variants (Extended Data Fig. 4c-e). This suggests that progenitors within a sub-lineage are at least partly coupled in their contribution to semen, likely due to non-random localization within or across the testes. Finally, in agreement with the absence of detectable selection, we noted no consistent linear changes, but rather random variation around a mean (Fig. 2f).

### Age-dependent changes observed in blood-specific but not sperm-specific variants

In the 5 AA individuals that underwent 300×MSMF we detected 120 *Sperm, 55 Shared*, and 1087 *Blood* mSNV/INDELs (Fig. 3a, Extended Data Fig. 5, Supplementary Dataset 1). AA individuals harbored a greater burden of *Blood* variants compared to YA; and two in particular, ID14 and ID17, had a further 5-fold increase, consistent with age-dependent ‘clonal hematopoiesis’^20,21^ (Fig. 3b). This phenomenon represents a loss of clonal diversity in the population of white blood cells as a result of selection or random drift with age^22^. *Shared* and *Sperm* variants, however, showed unchanged total numbers and mean AF across the age groups (Fig. 3c, Extended Data Fig. 6a-b), suggesting relative stability during aging.

**Figure 3.**
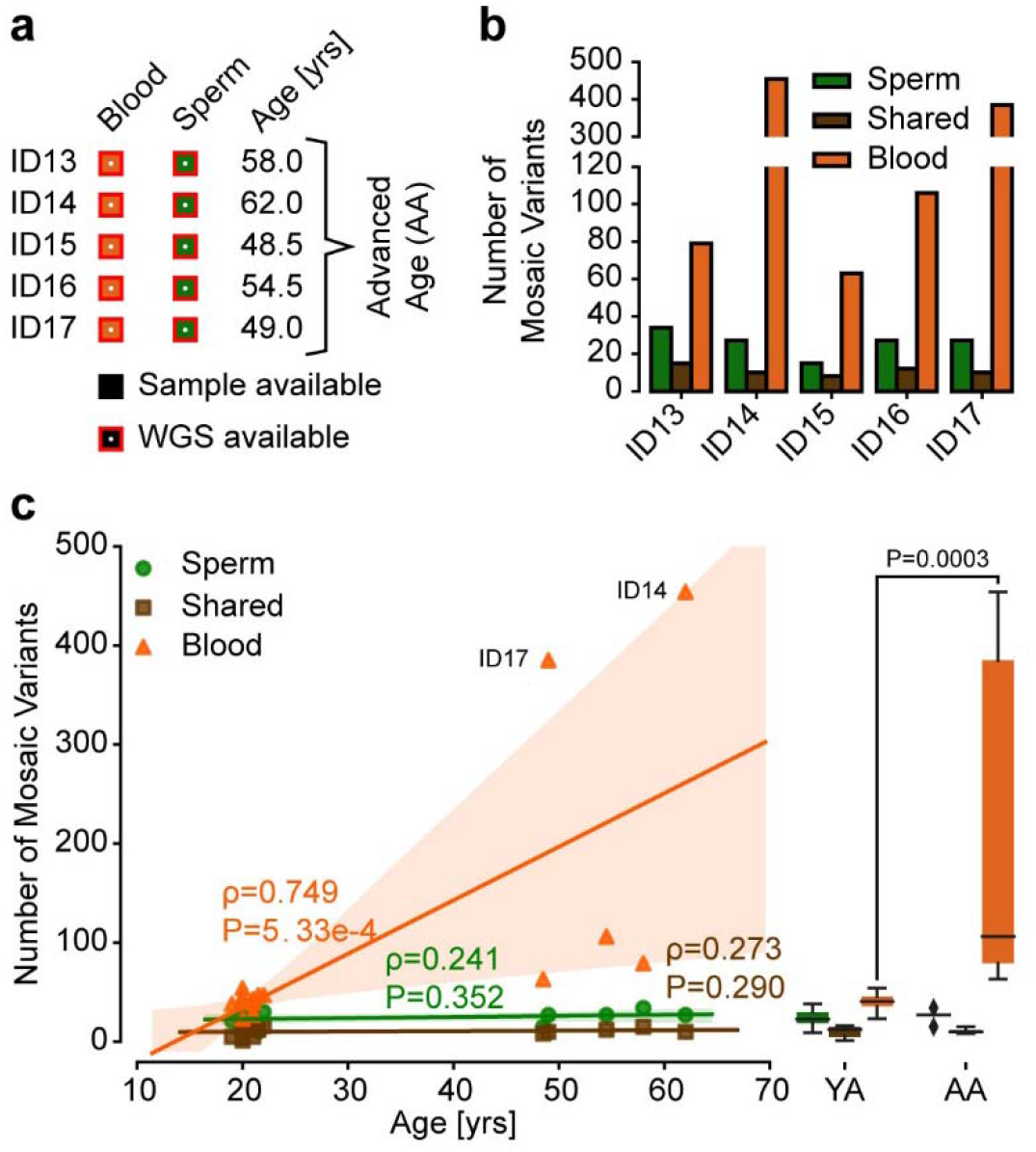
*Blood* but not *Sperm* variants increase with age. **a**, Available samples and ages for AA cohort individuals. **b**, Number of variants detected in the 5 AA individuals. Sperm mosaic variants were comparable to the YA cohort, but *Blood* variants were increased. **c**, Combined analysis of the number of YA and AA mosaic variants relative to the age of individuals. Left panel shows the data points, a regression line, and its 95% CI. Right panel shows a combined boxplot of all data points. For each of the three mosaic classes a two-tailed Mann-Whitney test was performed (Mann-Whitney U: *Sperm* 23, *Shared* 29.5, *Blood* 0; P-value: *Sperm* 0.4866, *Shared* 0.9764, *Blood* 0.0003).

Consistent with a positive selection for new somatic clones and their variants in hematopoietic lineages, we found a shift of *Blood* AFs towards lower abundance with age (Extended Data Fig. 6c-d). Unexpectedly, this was independent of whether the number of *Blood* variants was slightly (ID13, ID15, ID16) or greatly (ID14, ID17) increased, suggesting that detectable changes in blood mosaicism precede the collapse of clonal diversity, and that these changes can be identified in with 300xMSMF as early as the 5^th^ or 6^th^ decade.

### Mutational mechanisms change during early development

To increase the number of mosaic mutations available for aggregate analysis, we incorporated 200× deep WGS of sperm and blood from a previous study of 8 different men (REACH)^7,23,24^ and applied joint MSMF analysis. We found similar mosaicism patterns as in the YA and AA cohorts (Extended Data Fig. 7), and thus combined the previous 8 person and 17 person cohorts, yielding 522 *Sperm* and 251 *Shared* variants. In contrast, due to the clonal hematopoiesis detected in older men, we split blood-specific mosaicism into 473 ‘*Blood-Y*’ (YA) and 1673 ‘*Blood-A*’ (AA, REACH) variants (Fig. 4a). Of note, the last class was heavily biased towards three aged individuals with observed clonal hematopoiesis (Supplementary Dataset 1). These four aggregated classes were then used for a combined analysis of mutation features.

**Figure 4.**
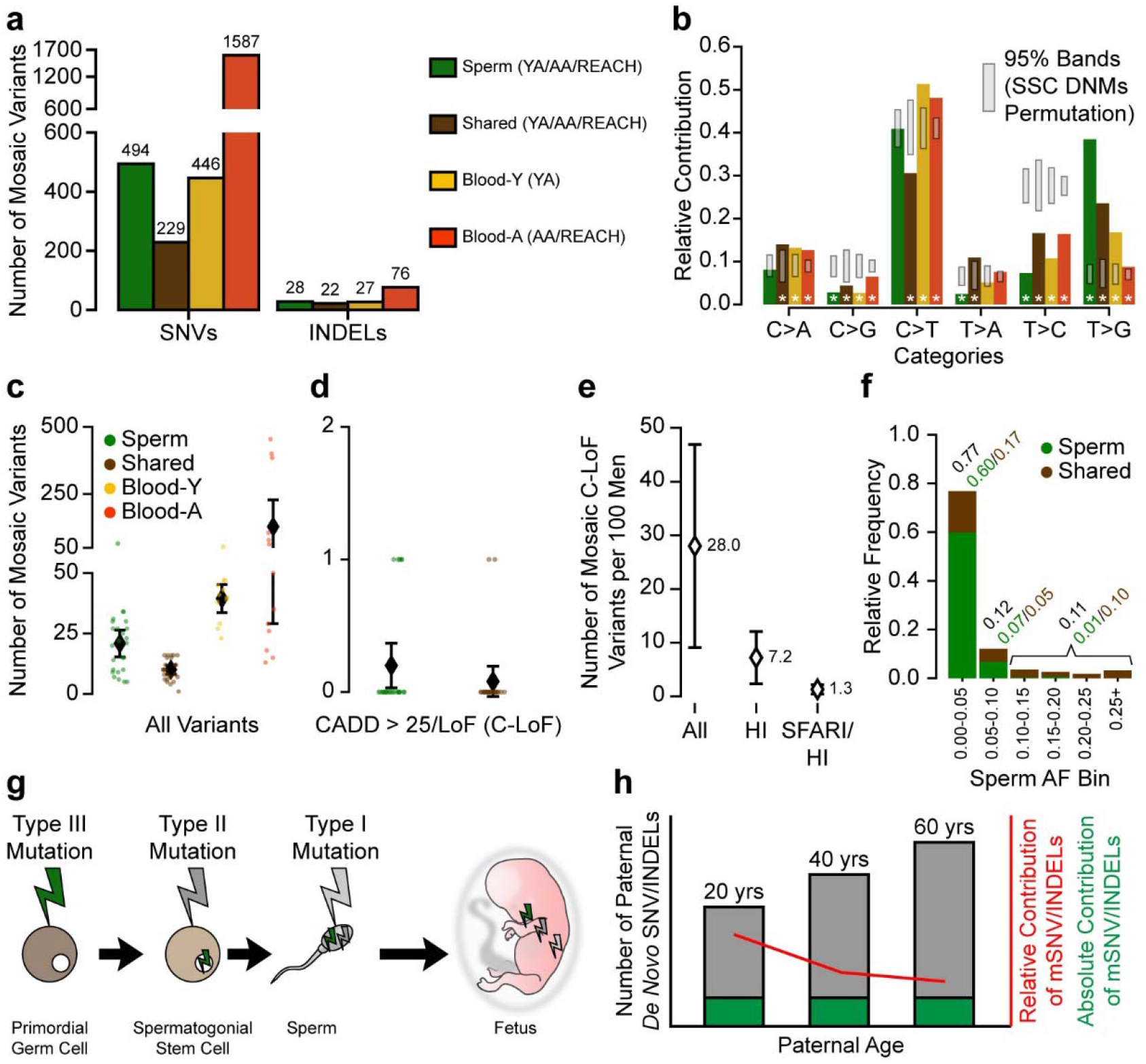
Sperm mosaicism accounts for a life-long transmission risk, by which approximately 7 in 100 males carry a high-impact, likely pathogenic mutation. **a**, Combination of the YA, AA, and REACH cohorts based on variant classes. Due to the previous analysis, *Blood* variants were split into *Blood-Y* from YA and *Blood-A* from AA and REACH, whereas *Sperm* and *Shared* variants were combined across cohorts. **b**, Single nucleotide substitution profiles of variant groups from a. Grey bands: 95% permutation intervals calculated from 10,000 random permutations of Simons Simplex Consortium *de novo* mutations (DNMs) from healthy siblings as baseline. Mosaic variants differed from permutations in several categories (asterisks). **c** and **d**, Detectable mosaic variants in each category for all variants (c) and *Sperm* and *Shared* for variants with a CADD score >25 or a loss-of-function prediction (d, C-LoF). Shown are data from each man, with mean with 95% confidence intervals. **e**, Estimated number of males per 100 (with 95% CI) with a detectable C-LoF variant (All), in a haploinsufficient (HI) gene, or in a haploinsufficient gene in the SFARI gene list (SFARI/HI). **f**, Relative frequency of AF categories, binned by 5% or above 25% for *Sperm* and *Shared* variants. The majority of mutations are <5% AF; however, most were not shared with blood. **g** and **h**, Three types of sperm mosaic variants. Type III are acquired during embryogenesis of the man, Type II accumulate during aging in spermatogonial stem cells, and Type I are acquired during or after meiosis. (g). While absolute contributions of Type III mutations are stable as men age, their relative contribution drops due to an age-dependent accumulation of other mutation types (h).

First, we contrasted substitution patterns of these four classes with matched permutations of variants from control germline mutations from the Simons Simplex Consortium^3^ or gnomAD^25^ (Fig. 4b, Extended Data Fig. 8a-e, Methods). The four mosaic classes showed distinct mutational patters from controls. For instance, C>G and T>C mutations were depleted among mosaic variants. *Shared* variants additionally had higher levels of T>A and lower levels of C>T mutations, thought to result from oxidative deamination^26^. The latter suggests that the impact of this mutational origin is relatively reduced in early embryonic development and replaced by others.

Next, we assessed mutation enrichment within genomic features compared to matched permutations (Extended Data Fig. 8f; Supplementary Dataset 3; Methods). *Blood-A* showed the most relative depletions, from areas of active histone modifications, annotated genes, low nucleosome occupancy, and sites associated with early replication timing, possibly indicative of selective pressure on clonal hematopoiesis clones. *Shared* variants were enriched in areas associated with late replication timing. *Sperm* variants showed enrichment in transcription factor binding sites, and a depletion in areas bound by Topoisomerases. In addition, both *Sperm* and *Blood-Y* variants were increased in DNase I hypersensitive sites, suggesting that open chromatin is more mutable in these two lineages.

Together, *Sperm* and *Shared* variants comprised more than 700 sperm mosaic mSNV/INDELs across the 25 individuals with a long tail of low AF mutations that were predominantly sperm-specific (Extended Data Fig. 8f-g). While mutations mainly accumulate as a function of cell cycle, some have suggested that this process is accelerated in less differentiated or early cells^27,28^, which would correlate with higher AFs. To assess this, we developed a quantitative metric termed ‘Mutation Factor’, which is defined by the rate of mutational accumulation as a function of predicted developmental ‘age’. We determined this metric by fitting a step-wise exponential regression with minimal loss to the ranked plot of mosaic variants (Extended Data Fig. 9a-e, Methods). We found almost identical Mutation Factors for *Shared* variants in blood and sperm, suggesting transmission to both tissues at stable fractions. *Sperm* and *Blood-Y* variants also had a comparable Mutation Factor, supporting a similar accumulation of mutations that was more dependent on the developmental time rather than the fate of the progenitors. *Shared* variants accumulated at a faster rate per cell cycle than *Sperm* or *Blood-Y* variants, as was postulated to result from DNA damage repair differences in early development^27–29^. *Blood-A*, however, had an even higher Mutation Factor than the other classes, likely reflecting of the dynamic changes in clonal proportions with aging. These observations were largely confirmed by quantile analysis of AF distributions (Extended Data Fig. 9f).

### One in 15 men harbors a likely pathogenic transmissible mutation in sperm

Across all cohorts, men harbored an average of 30.9 sperm mosaic variants (*Sperm*: 20.9, *Shared*: 10.0) (Fig. 4c). Of these, 1.6 (*Sperm*: 1.1, *Shared*: 0.5) were exonic (Extended Data fig. 10a), and 0.3 (*Sperm*: 0.2, *Shared*: 0.1) of ‘high-impact’, i.e. CADD^30^ score—a metric that summarizes deleteriousness of a mutation—above 25 or predicted loss-of-function (C-LoF; Fig. 4d, Supplementary Dataset 3). Thus, across 100 men, 28 are predicted to harbor a C-LoF variant in sperm at detectable AFs, and 7.2 in a haploinsufficient (HI) gene^31^ (Fig. 4e), amounting to approximately 1 in 15 males; and—based on our observations in the YA and AA cohort—this risk should be stable for life. The intersection of these genes with the resultant disease in the setting of a mutation could define the disease-specific transmissible burden for conditions with near-Mendelian risk. For instance, 1.3 in 100 men are estimated to carry a mutation with potential to cause monogenetic autism spectrum disorder (Fig. 4e). For genes with lower risk of disease when mutated, the odds ratio of the disease (i.e. risk of disease when the gene is mutated) need also be taken into account.

Overall, most variants in the combined cohort were at AFs between 1-26% (Extended Data Fig. 10b), with the majority of sperm AFs—and thus likely transmission risks—below 5% (Fig. 4f), suggesting that the risk of transmission for any given variant is relatively low. But approximately 1 in 5 variants had a higher AFs, and the majority of these were detectable in both blood and sperm. Adjusted for relative frequency and AF, *Sperm* and *Shared* variants represented a similar total transmissible burden. Assessment of sperm for mosaicism is critical to assess risk of transmission, because using blood as a surrogate will produce false-negatives as a result of sperm-specific variants and false-positives as a result of blood*-*specific variants, the latter increasing as a function of aging.

## Discussion

Here we provide an overview of the landscape of sperm mosaicism through comprehensive assessment using deep WGS of sperm and blood across multiple men, multiple time points, and multiple ages. AFs correlated with tissue specificity of mutations, with *Shared* variants showing early embryonic origins and higher AFs, compared with *Sperm* or *Blood* variants showing origins after lineage separation with lower AFs. PGCs separate from somatic progenitors before the third week post conception in humans^32^ (Extended Data Fig. 10d). Hematopoietic progenitors arise later from mesoderm, following germ layer specification^33^. Thus, *Blood* mutations detected in YA are—in contrast to *Sperm*—likely shared with other tissues. Our *Sperm* variant AFs estimate that the number of starting progenitors committed to the sperm lineage in development as 3-6 cells. It is difficult, however, to provide such an estimate for blood due to the changing nature of clonal composition over time.

*Shared* variants differed in their mutational accumulation speed from those that were tissue-specific, consistent with previous studies^27,28^, and where deficiencies in early repair mechanisms have been proposed as a possible explanation^29^. Supporting our idea that *Sperm* and *Blood* variants occur at a similar time in development, *Sperm* and *Blood-Y* mutations exhibit comparable mutational spectra and Mutation Factors. Due to the likely embryonic origin of detectable mosaic mutations, we observed a depletion of T>C variants that correlate with gonadal aging^5,6^. Thus, these mutations differ from those due to Type I or II mosaicism, which are sporadic or potentially accumulated with age, and which could be directly assessed with future single cell sperm sequencing studies.

Our analysis provides evidence that Type III sperm mosaic variants are stable in number and AF, likely across the entire lifetime of an individual. Thus, Type III variants represent a persistent, long-term transmission risk (Extended Data Fig. 10e). However, their relative contribution to disease risk actually declines with age, due to the increase of Type I and II mutations (Fig. 4g-h). These ideas are consistent with population analysis and recurrence risk estimates that consider age^6,14^. Consequently, spermatogonial stem cells, despite proliferating throughout reproductive life, do not appear to exhibit detectable clonal collapse or expansion, likely a reflection of cellular dynamics in the testicular stem cell niche. Single cell sequencing identified almost 3% of sperm with karyotype defects^34^, but the degree to which they contribute to risk of disease in offspring remains unknown.

Previous work has demonstrated risk of clonal hematopoiesis with advancing age^20,21,35^. We provide genome-wide analysis documenting evidence for this phenomenon in individuals as young as 40s-50s, the earliest to date. Whereas *Blood* variants were substantially elevated in only some AA individuals, all AAs exhibited changes in AF spectrum compared with YA. While the origins of clonal hematopoiesis remain controversial^22^, there appear to be selective pressures during hematopoiesis in most or all aging individuals that do not extend to their gonadal lineage (Extended Data Fig. 10e).

Sporadic mutations are major contributors to congenital human disease^4,36,37^, compounded by increased parental age that correlates with *de novo* mutation burden^1,5^. While we do not address this problem directly, we provide a framework by which Type III mosaicism could be detected for genetic counselling decisions. We predict that approximately 1 in 300 concepti harbors a variant that is likely pathogenic, causing miscarriage or congenital disease. As a consequence, for the monogenetic component of a well-studied disorder like autism, we estimate that ~15% are caused by transmitted Type III mosaicism (see Methods). We show that natural variation of sperm mosaicism in an individual is modest over time, and that these variants have their origin when the father (or future father) was an embryo himself. Our approach could be applied to formulate effective prevention strategies for potentially harmful mutations.

## Acknowledgements

We thank the participants in this study for their contribution. We also thank Drs. Yong E. Zhang, Johannes C. M. Schlachetzki, and Alan Packer for feedback and suggestions. M.W.B. was supported by an EMBO Long-Term Fellowship (ALTF 174-2015), which is co-funded by the Marie Curie Actions of the European Commission (LTFCOFUND2013, GA-2013-609409), and an Erwin Schrödinger Fellowship by the Austrian Science Fund (FWF): J 4197-B30. This study was supported by grants to J.G.G. from the NIH (U01MH108898, R01NS083823), and the Simons Foundation Autism Research Initiative (SFARI, 571583). Sequencing support was provided by the Rady Children’s Institute for Genomic Medicine. We thank R. Sinkovits, A. Majumdar, S. Strande, and the San Diego Supercomputer Center for hosting the computing infrastructure necessary for completing this project.

## Author Contributions

M.W.B, X.Y., and J.G.G. conceived the project and designed the experiments. X.Y., M.W.B., L.L.B., S.A.W., and G.C. performed the experiments. X.Y., X.X., M.W.B., D.A., R.D.G., B.C., and A.N. performed the bioinformatic and data analyses. D.A. and J.S. performed the analyses of the REACH cohort. K.N.J., V.S., J.M.-V., and M.W.B. requested, organized, and handled patient samples. S.N., L.V.D.K., and YD performed sequencing analysis. M.W.B., X.Y., and J.G.G. wrote the manuscript. J.G.G. supervised the overall project. All authors have seen and commented on the manuscript prior to submission.

## Methods

### Subject recruitment

17 healthy males were enrolled according to approved human subjects protocols from the Institutional Review Board (IRB) of the University of California for blood, saliva, and semen sampling (140028, 161115). All participants signed informed consents according to the IRB requirement, and the study was performed in accordance with Health Insurance Portability and Accountability Act (HIPAA) Privacy Rules. None of the participants reported severe psychological conditions or showed significant signs of neurological disorders, infectious diseases, or cancer. Semen and blood samples were collected for all subjects (ID01-17). ID01-08 and ID11 further provided saliva samples. ID05 further provided a second semen sample approximately half a year after first collection; ID02-04, ID06-S08, ID11, and ID12 provided a total of 3 samples within ~12 months.

### DNA extraction for blood and saliva

Genomic DNA was extracted from peripheral blood and saliva samples containing buccal cells using the Puregene kit (Qiagen, #158389) following the manufacturers’ recommendations.

### Sperm extraction

Extraction of sperm cell DNA from fresh ejaculates was performed as previously described^7,38^. In short, sperm cells were isolated by centrifugation of the fresh (up to 2 days) ejaculate over an isotonic solution (90%) (Sage/Origio, ART-2100; Sage/Origio, ART-1006) using up to 2 mL of the sample. Following a washing step, quantity and quality were assessed using a cell counting chamber (Sigma-Aldrich, BR717805-1EA). Cells were pelleted and lysis was performed by addition of RLT lysis buffer (Qiagen, 79216), Bond-Breaker TCEP solution (Pierce, 77720), and 0.2 mm stainless steel beads (Next Advance, SSB02) on a Disruptor Genie (Scientific Industries, SI-238I). The lysate was processed using reagents and columns from an AllPrep DNA/RNA Mini Kit (Qiagen, 80204). Concentration of the final eluate was assessed employing standard methods. Concentrations ranged from ~0.5-300 ng/μl.

### WGS of sperm and blood samples

WGS sequencing was performed as described before^18^. In short, a total of 1.0 μg of extracted DNA was used as the starting material for PCR-free library construction (KAPA HyperPrep PCR-Free Library Prep kit; Roche, #KK8505); libraries were then mechanically sheared (Covaris microtube system; Covaris, # SKU 520053) to obtain ~400 base pairs (bp) fragments. Then Illumina dual index adapter were ligated to these DNA fragments. Following beads-based double size selection (300-600 bp), the concentration of ligated fragments in each library was quantified (KAPA Library Quantification Kits for Illumina platforms; Roche/KAPA Biosystems, # KK4824). Libraries with concentrations of more than 3 nM and fragments with peak size 400 bp were sequenced on an Illumina Novaseq 6000 S4 and/or S2 Flow Cell (FC), in 6-8 independent pools. The target for whole genome sequencing with high quality sequencing raw data was 120 GB or greater with a Q30 >90% per library per sequencing run. In case the first sequencing run generated less than that, additional sequencing was performed by sequencing the same library on a Novaseq 6000 S2 FC with 2×101 read length. Raw data was processed through the DRAGEN platform to generate BAM files.

### WGS data processing and germline variant calling

Raw data was aligned to the GRCh37d5 reference genome, sorted, and PCR duplicates were removed by a DRAGEN platform. Reads aligned to the INDEL regions were realigned with GATK’s (v3.8.1) RealignerTargetCreator and IndelRealigner following the GATK best practice. Base quality scores were recalibrated using GATK’s (v3.8.1) BaseRecalibrator and PrintReads. Read groups were renamed by Picard’s (v2.20.7) AddOrReplaceReadGroups command. Germline SNVs and INDELs were detected by GATK’s (v3.8.1) HaplotypeCaller. The distribution of library DNA insertion was assessed by Picards’ (v2.20.7) CollectInsertSizeMetrics. The depth of coverage was analyzed by GATK’s (v3.8.1) DepthOfCoverage command.

### Principle component analysis (PCA) of genetic origins of the assessed individuals

In order to determine the origins of the included individuals, heterozygous variants generated by GATK’s (v3.8.1) HaplotypeCaller were output in genomic VCF format and genotyped across samples by using the GATK’s (v3.8.1)’s GenotypeGVCFs and CombineGVCFs; in addition all variants from dbSNP (v137) were added. The VCF file was reformated by BCFtools (v1.10.32) and converted to bfiles by PLINK (v1.90b6.16). Single nucleotide polymorphisms (SNPs) were extracted from both the samples in this study and samples from the 1000 Genomes phase 3^39^ and merged together; any SNP that overlapped with the repeat mask region was removed. PCA was carried out by PLINK (v1.90b6.16) and the results were plotted in R (v3.5.1).

### Mosaic SNV/INDEL detection pipeline in WGS data (300×MSMF)

Mosaic single nucleotide variants/mosaic small (typically below 20 bp) INDELs were called by using a combination of four different computational methods based on previous published and adapted pipelines^7,18^: the intersection of variants from the paired-mode of GATK’s (v4.0.4) Mutect2^40^ (paired mode) and Strelka2^41^ (v 2.9.2) (set on ‘pass’ for all variant filter criteria) for sample-specific variants; or single-mode of Mutect2 (with an in-house panel of normal) followed by MosaicForecast^17^ (v0.0.1) for sample-specific or tissue-shared variants. For the YA cohort, the panel of normal is generated using a “leave one out” strategy, by excluding samples from each individual; for the AA and REACH cohort, all samples from the YA were used to generate the panel of normal. Variants were excluded if they 1] resided in segmental duplication regions as annotated in the UCSC genome browser (UCSC SegDup) or RepeatMasker regions, 2] resided within a homopolymer or dinucleotide repeat with more than 3 units, 3] overlapped with annotated germline INDELs, 4] did not show a minimum of 3 alternative reads, or 5] were detected more than once across multiple individuals. We further removed any variants with a population allele frequency (AF) >0.001 in gnomAD^25^ (v 2.1.1) or >0 for variants only detected by MosaicForecast^17^. To avoid binomial sampling bias and false positive signal from copy number/structural variations or non-annotated repetitive regions, we randomly chose 1600 single nucleotide polymorphism from dbSNP (v137), estimated the 95% confidence interval of all those variants in each sample respectively, and excluded variants whose coverage is not within this CI. Finally, variants with an AF>0.35 in both sperm and blood (or >0.7 for sex chromosomes) were considered likely germline variants and removed. Variants with a lower CI of AF<0.001 were also removed. Fractions of mutant alleles for variants called in one sample were calculated in the other sample with the exact binomial confidence intervals using scripts described below. If a variant was only detected in one tissue, mosaicism in the second tissue was confirmed if a minimum of 3 alternative reads were present. Scripts for variant filtering and annotations are provided on GitHub (https://github.com/shishenyxx/Sperm_control_cohort_mosaicism).

### Simulation analysis to determine the sensitivity of 300×MSMF

To determine the sensitivity for detecting mosaic variants, we created simulated datasets that contained known mosaic variants at low frequencies. We first randomly generated 10,000 variants from chromosome 22 based on GRCh37d5 as our set of mosaic variants. We then used Pysim^42^ to simulate Illumina paired-end sequencing reads with a NovaSeq 6000 error model from the GRCh37d5 reference chromosome 22 and a version of chromosome 22 that contained the alternate alleles from our 10,000 mosaic variants. These two sets of reads were then combined to create a series of datasets with mosaic variants at 1,2, 3, 4, 5, 10, 15, 20, 25, and 50% AF, at coverages at 50×, 100×, 200×, 300×. 400×, and 500× depth. Reads were mapped to GRCh37d5 using BWA (v0.7.8) mem, processed with Picard’s (v2.20.7) MarkDuplicates, and INDELs were realigned and base quality scores recalibrated as described above. We applied our somatic variant calling pipelines containing GATK’s (v4.0.4) Mutect2 (single mode and paired mode), Strelka2 (v 2.9.2), and MosaicForecast (v0.0.1) to detect mosaic variants at each AF and each depth. We further applied the same filters we used for the genomic regions; as we excluded the repetitive and segmental duplication regions, only 75% of the genomic region remained valid. The sensitivity and recovery rate of the pipeline was then determined through these data.

### Visualization of genomic distribution of mosaic variants

The genomic distribution pattern of mosaic variants and the allelic fractions of different variants across the genome was presented using Circos^43^ (v0.69-6).

### Targeted amplicon sequencing (TAS) and experimental benchmark of the SNV/INDEL calling pipeline

TAS analysis was first applied to 185 variants from the previously published 200× WGS sequencing results^7^ to experimentally confirm the validation rate of the new pipeline. PCR products for sequencing were designed with a target length of 160-190 bp with primers being at least 60 bp away from the base of interest. Primers were designed using the command-line tool of Primer3^44,45^ with a Python (v3.7.3) wrapper^7^. PCR was performed according to standard procedures using GoTaq Colorless Master Mix (Promega, M7832) on sperm, blood, and an unrelated control. Amplicons were enzymatically cleaned with ExoI (NEB, M0293S) and SAP (NEB, M0371S) treatment. Following normalization with the Qubit HS Kit (ThermFisher Scientific, Q33231), amplification products were processed according to the manufacturer’s protocol with AMPure XP Beads (Beckman Coulter, A63882) at a ratio of 1.2x. Library preparation was performed according to the manufacturer’s protocol using a Kapa Hyper Prep Kit (Kapa Biosystems, KK8501) and barcoded independently with unique dual indexes (IDT for Illumina, 20022370). The libraries were sequenced on an Illumina HiSeq 4000 platform with 100 bp paired-end reads. After determining the validation rate of the new pipeline, TAS was further performed for a subset of called variants on the different sperm time points, blood, saliva, and unrelated control sample to quantify the AFs and to extend analysis to tissues that were not subjected to WGS.

### Data analysis for TAS

Reads from TAS were mapped to the GRCH37d5 reference genome by BWA mem and processed according to GATK (v3.8.2) best practices without removing PCR duplicates. Putative mosaic sites were retrieved using SAMtools (v1.9) mpileup and pileup filtering scripts described in previous TAS pipelines^7^. Variants were considered mosaic if 1] their lower 95% exact binomial CI boundary was above the upper 95% CI boundary of the control; 2] their AF was >0.5%. Sperm samples from the YA cohort were labeled as time point 1 (t_1_), t_2_ and t_3_, based on the data of sample collection. t_1_ was used as an anchor to determine absolute and relative (i.e. fold change) AF differences of the same variant measured across samples.

### Mutational signature analysis

Mutational signatures were determined for each variant by retrieving the tri-nucleotide sequence context using Python (v3.5.4) with pysam (v0.11.2.2) and plotting the trans- or conversion based on the pyrimidine base of the original pair similar to previous studies^46^. Mutational signatures from *de novo* mutations in the Simons Simplex Consortium cohort (from healthy siblings) and general mutations from gnomAD were obtained by retrieving SNVs present in their respective, publicly available VCFs. In order to obtain a 95% band of expectation, an equivalent number of variants was randomly chosen from the Simons Simplex Consortium or gnomAD VCF. This process was performed for a total of 10,000 times to obtain a distribution and the 2.5th and 97.5th percentile of the simulated mutational signatures. Significance was reported if a mutational signature was outside the permuted 95% bands.

### Step-wise exponential regression model for the burden of variants

In order to model the exponential decay of the variants, a step-wise exponential regression model was made based on the following assumptions: 1] variants happening at roughly the same cell division during early embryonic development have similar allelic fractions in different individuals; 2] during early embryonic development the number of cells are growing exponentially but at different rates across tissues due to varying growth rates and cell death; 3] the spontaneous mutation rate is stable within each category; 4] the number of mosaic variants occurring in each cell generation is in proportion with the number of cells in that generation. For each group of ranked variants from an already developed tissue (sperm or blood), during the *t^th^* cell division, we assume that all variants came from a starting population of 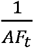 variants, and *AF*_0_ is estimated from the exact binomial CI of the highest AFs found in each group. Based on assumption 2 the mutation is accumulated at a speed of *θ* (*θ* ≥ 1 and *θ* ≤ 2). For the *t^th^* cell division, the average 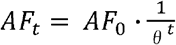, and the number of expected variants with this *AF_t_* is 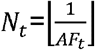, we rank the *AF_t_* to get an estimated rank vector 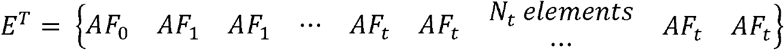, to get a best estimation of *E^T^* towards the observed ranked AF vector *O^T^*, we defined the loss 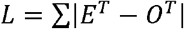. By minimizing *L*, we obtained the best estimation of the ranked AF curve. We finally defined a 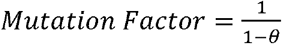 as the output to reflect the mutation burden from embryonic development to the time of sampling. The higher the mutation burden, the higher the mutation factor, in order to reach the accumulated number of ranked variants.

### Assessment of mosaic variants overlap with different genomic features

In order to assess the distribution of mosaic variants and their overlap with genomic features, an equal number of variants (mSNV/INDELs that were *Sperm, Shared, Blood-Y*, and *Blood-A*) was randomly generated with the BEDTools (v2.27.1) shuffle command within the region from Strelka 2 without the subtracted regions (e.g. repeat regions). This process was repeated 10,000 times to generate a distribution and their 95% CI. Observed and randomly subsampled variants were annotated with whole-genome histone modifications data for H3k27ac, H3k27me3, H3k4me1, and H3k4me3 from ENCODE v3 downloaded from the UCSC genome browser (http://hgdownload.soe.ucsc.edu/goldenPath/hg19/database/)—specifically for the overlap with peaks called from the H1 human embryonic cell line (H1), as well as peaks merged from 10 different cell lines (Mrg; Gm12878, H1, Hmec, Hsmm, Huvec, K562, Nha, Nhek, and Nhlf). Gene region, intronic, and exonic regions from NCBI RefSeqGene (http://hgdownload.soe.ucsc.edu/goldenPath/hg19/database/refGene.txt.gz); 10 Topoisomerase 2A/2B (Top2a/b) sensitive regions from ChIP-seq data^47^ (Samples: GSM2635602, GSM2635603, GSM2635606, and GSM2635607); CpG islands: data from the UCSC genome browser (http://hgdownload.soe.ucsc.edu/goldenPath/hg19/database/); genomic regions with annotated early and late replication timing^48^; nucleosome occupancy tendency (high/>0.7 or low/0.0-0.7 as defined in the source) from GM12878, for which all non-zero values were extracted and merged^49^; enhancer genomic regions from the VISTA Enhancer Browser (https://enhancer.lbl.gov/); and DNase I hypersensitive regions and transcription factor binding sites from Encode v3 tracks from the UCSC genome browser (wgEncodeRegDnaseClusteredV3 and wgEncodeRegTfbsClusteredV3, respectively).

### Prediction of loss-of-function

Variants were annotated as loss-of-function if both the SVM-based iFish^50^ probability equaled 1 and the DeepLearning-based Define^51^ was >0.7, or if it was annotated as frameshift with a gnomAD allele frequency <0.0001.

### Burden Estimation

Using the observed fraction of variants that are classified as C-LoF, we calculated a 95% estimation interval of the true fraction using SciPy (v1.3.1) stats’s t-interval and multiplied by the chosen number of men (n=100). This fraction was further modified by taking into account the subset of genic regions that are annotated to belong to a haploinsufficent gene (HI) by the definition of ClinGen (https://dosage.clinicalgenome.org/help.shtml) level 3 “Sufficient evidence for dosage pathogenicity” or that belong to an HI gene which is annotated as a likely autism spectrum disorder gene by SFARI (Level 1,2,3, and S, with PLI higher than 0.9). Genomic regions of those genes were extracted from http://www.openbioinformatics.org/annovar/download/hg19_refGene.txt.gz.

### Estimation of disease impact conveyed by Type III mosaicism

For transmission risk we assume that 1] expression of the disrupted gene does not impact a sperm cell’s fertility; 2] AFs estimated in purified sperm directly reflect the percentage of sperm cells carrying the mutation and determine the average transmission risk *θ*. For any disease with incident rate *I* and a fraction *P*, which are caused by *de novo* HI-C-LoF SNV/INDELs within a set of genes *HI – C – LoF ⋂ Disease gene set* (monogenetic, autosomal dominant contribution), we can calculate the percentage of the relevant genome by comparing 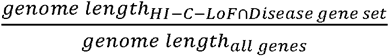. Taking *μ* into account, which is the fraction of men predicted to carry a C-LoF mutation, we can estimate the explained risk for a specific disease/phenotype with

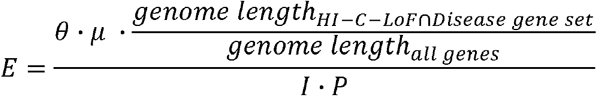

Taking ASD as an example, exonic *de novo* C-LoF SNV/INDELs contribute to *P* = 21% of ASD diagnose^3^. According to the CDC, in 2020, approximately *I* = 1/54 children in the US is diagnosed with ASD (https://www.cdc.gov/ncbddd/autism/data.html). Roughly *I · P* = 3.89/1000 children are born with ASD caused by *de novo* C-LoF SNV/INDELs. Our data determines an average *θ* = 0.047 and a *μ* = 0.27, and thus 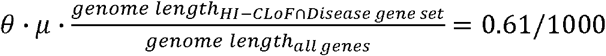, assuming that ASD HI-C-LoF mutations do not increase miscarriage rates. Therefore, Type III mosaicism described in this manuscript contributes an estimated *E* ≈ 1/6 of *de novo* SNV/INDELs underlying ASD diagnose. As those mutations are of early embryonic origin, prior to sex divergence, this contribution should be similar in both parents^52^, suggesting that overall, parental gonadal mosaicism contributes 1/3 of *de novo* ASD SNV/INDELs. This approach can be easily extended to other diseases or phenotypes with known monogenetic architecture, such as epilepsy, intellectual disabilities, or congenital heart disease^53,54^. Note that the HI-C-LoF themselves, based on the data and considerations outlined above, will be transmitted to ~1 in 300 concepti, likely leading to miscarriage or congenital disease.

### Data processing

Data analysis and plotting were performed using R (v3.5.1) with ggplot2 (v3.3.1) and Rcpp (v1.0.3) packages; or with Python (v3.6.8) with pandas (v0.24.2), matplotlib (v3.1.1), numpy (v1.16.2) SciPy (v1.3.1) and seaborn (v0.9.0) packages.

### Statistical analyses

Statistical analyses were performed with R (Spearman, exact binomial confidence intervals, quantile analysis, and Kolmogorov-Smirnov test), GraphPad Prism (Mann-Whitney Test), and Python with pandas (95% confidence interval determination).

### Data availability statement

Aligned BAM files generated for this study through deep WGS or TAS are available on SRA (accession number: PRJNA660493 and PRJNA588332). Data are also available through the corresponding authors on reasonable request. Additionally, summary tables of the data are included as supplementary information.

### Code availability statement

Codes for data analysis pipelines as well as codes to generate the figures are freely available on github at https://github.com/shishenyxx/Sperm_control_cohort_mosaicism.

## Extended Data Figures

**Extended Data Figure 1.**
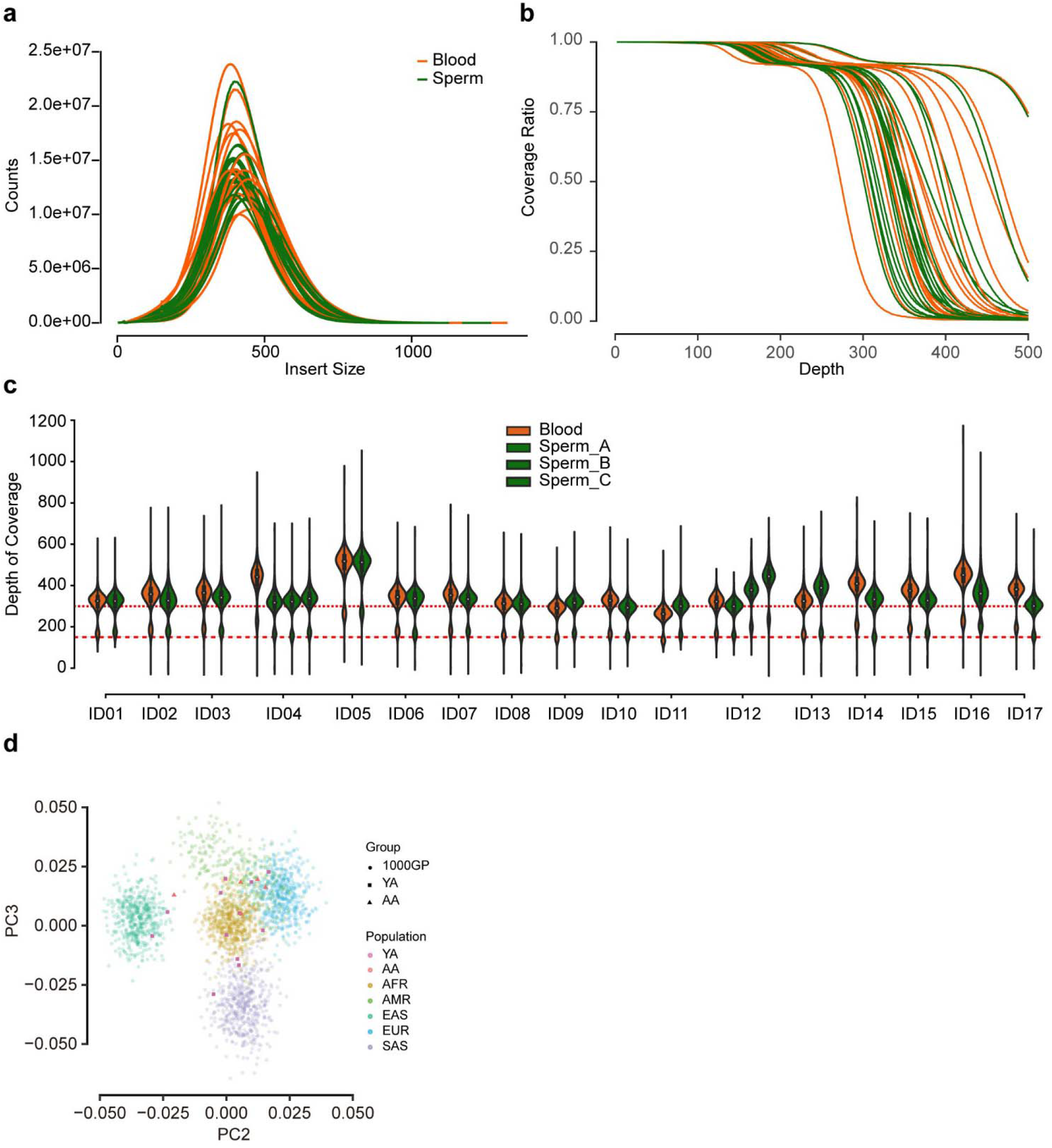
Quality control for WGS data. **a**, Library DNA insert size for each whole-genome sequencing (WGS) sample. All the samples had a consistent single peak at ~400 base pairs. Colors were used to distinguish different sample types. **b**, Cumulative proportion of coverage depth of each WGS sample. The majority of samples reached 300×, two samples from one individual reached 500×. **c**, Distribution of depth of 1600 SNVs (randomly selected from dbSNP v137) detected in each WGS sample. Almost all variants from the autosomes reached 300× and variants from sex chromosome reached 150×, 95% confidence of the depth of autosome and sex chromosome were calculated to exclude extreme depth from copy number (CNVs) or structural variants (SVs). **d**, Principal component analysis revealing ancestry of the donors for the study; compared with the data from the 1000 Genome project phase3 release, the individuals from the YA and AA cohorts showed a variety of ancestries. 1000GP: individuals from the 1000 Genome phase3 collection; AFR: African; AMR: Admixed American; EAS: East Asian; EUR: European; SAS: South Asian

**Extended Data Figure 2.**
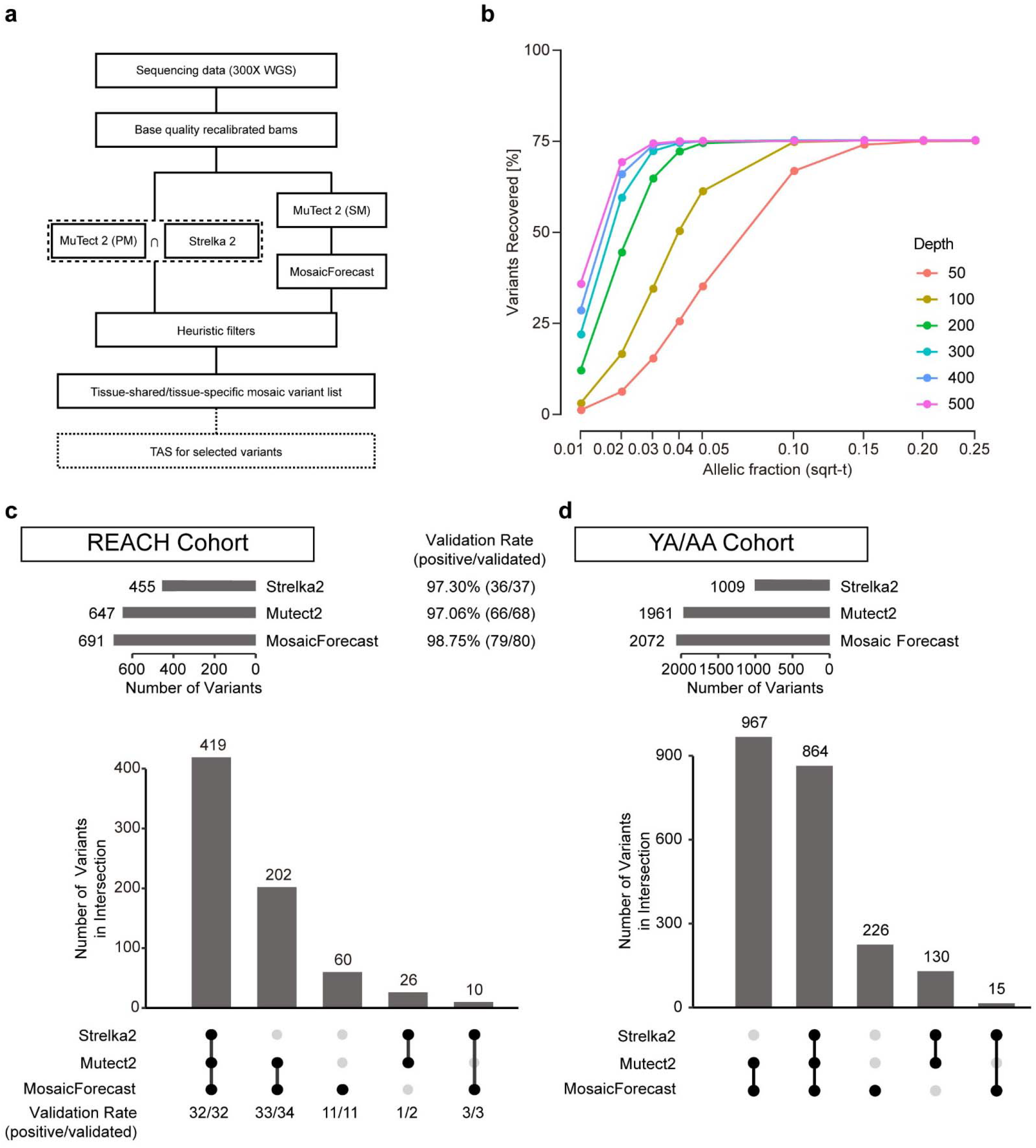
Computational and experimental benchmarks of the analytical workflow showed high validation rates. **a**, Workflow for sequencing data processing, mosaic variant calling, and variant quantification. Details are described in the Methods section. The 300×MSMF pipeline is a combination of MuTect2, Strelka2, and the state-of-the-art MosaicForecast classifier. PM: paired mode, SM: single mode. TAS: targeted amplicon sequencing. **b**, Computational benchmark for the detection sensitivity of the workflow. The x-axis shows the expected AFs and y-axis the sensitivity; colors distinguish different simulated read depths. Among the 10,000 simulated variants, 75% fall in non-repetitive regions. The 25% repetitive regions were hard-filtered at the ‘heuristic filters’ step in our pipeline. For low-AF variants at 1-2%, 300-500× sequencing showed similar detection sensitivity, but significantly improved compared with 200×, 100×, and 50×. For high-AF variants >20%, all depths showed maximal sensitivity. **c**, UpSet plot showing reanalysis of data from the previously described REACH cohort^7^ using the described workflow. Plot shows the yield and TAS validation rate for each individual variant caller within the mosaicism detection workflow. Overall, the workflow has a 97.6 (80/82) validation rate. **d**, UpSet plot for the YA and AA cohort showed similar relative numbers of mosaic variants detected by each method, although MosaicForecast had a higher relative yield.

**Extended Data Figure 3.**
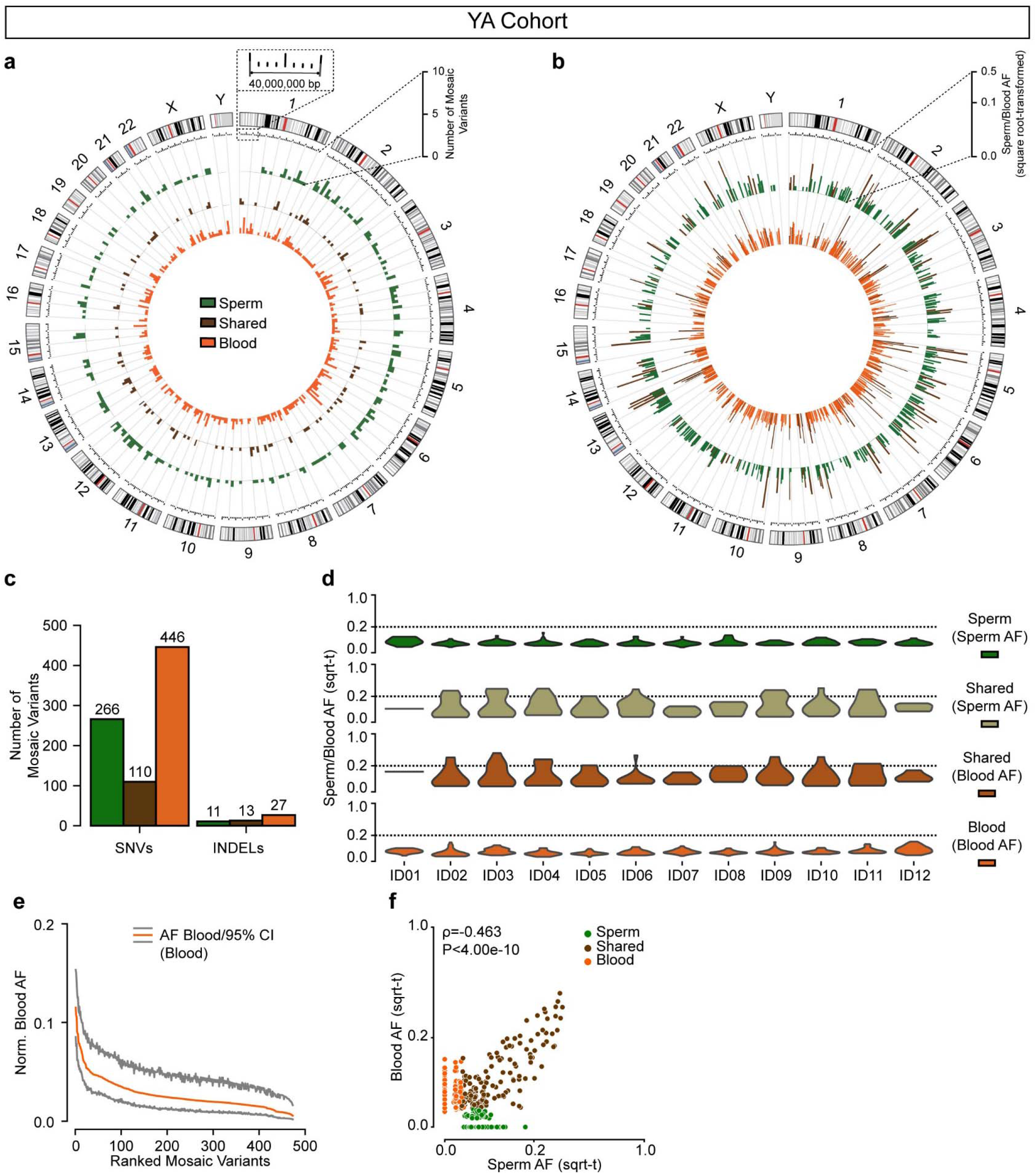
Genomic distribution and allelic fraction (AF) properties of variants detected in the YA cohort. **a**, Circos histograms for the number of mSNV/INDELs detected from the YA cohort, colors distinguish classes of the variants: green, sperm-specific (*Sperm*); brown: tissue-shared (*Shared*); orange: blood-specific (*Blood*); circles from inner to outer are number of mosaic variants detected only in blood (*Blood*), detected in both blood and sperm (*Shared*) and detected only in sperm (*Sperm*). Variants were evenly distributed across the genome. **b**, Mosaic SNV/INDELs and the corresponding allelic fractions (AFs) detected from the YA cohort, colors are the same as (a); Inner circle: AFs in the blood; outer circle: AFs in the sperm. **c**, Number of mosaic SNVs and INDELs detected from the YA cohort. Two-thirds of the transmissible variants were only detectable in sperm. **d**, AF distribution (square root-transformed; sqrt-t; for orientation 0.2 AF is highlighted with a dotted line) of *Sperm, Shared*, and *Blood* variants within each individual. *Shared* variants show higher peak and overall AF compared to *Sperm* and *Blood*. **e**, Ranked plot of the estimated blood AF with 95% confidence intervals of *Blood* variants. **f**, Correlation between the square root-transformed (sqrt-t) AFs from WGS in blood and sperm of the YA cohort for all detected variants.

**Extended Data Figure 4.**
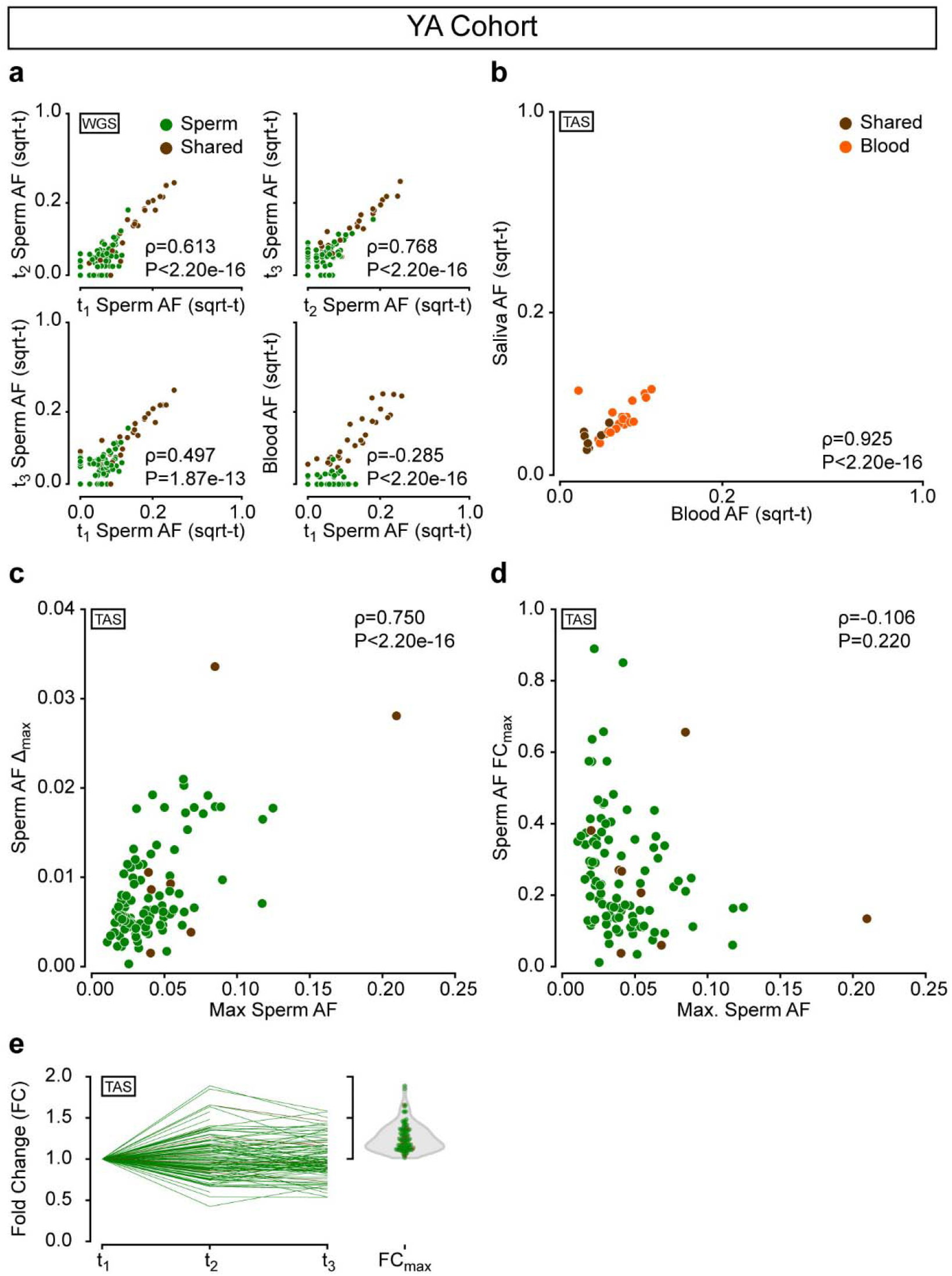
Correlation plots of AFs in different samples from the YA cohort. **a**, Pair-wise AF comparisons of sperm mosaic variants from ID04 and ID12 analyzed by WGS. Sperm samples showed high correlation, but some variants were only detectable in one sperm sample. **b**, Correlation between sqrt-t AFs from TAS in blood and saliva in YA. **c**, Correlation of maximum AF in any of the 3 sperm samples and the maximum differences between the 3 sperm AFs from TAS. The positive correlation suggests increased variation of absolute differences with increased AF. However, this correlation is imperfect. **d**, Correlation of maximum AF in any of the 3 sperm samples and the maximal fold change between the 3 sperm AFs from TAS. No significant correlation is observed. This suggests that smaller AF variants may experience outsized, relative variability. All panels show Spearman’s ρ and P-value across all variants. **e**, Fold change for each tested variant. Variation can reach up to 1.5-2-fold.

**Extended Data Figure 5.**
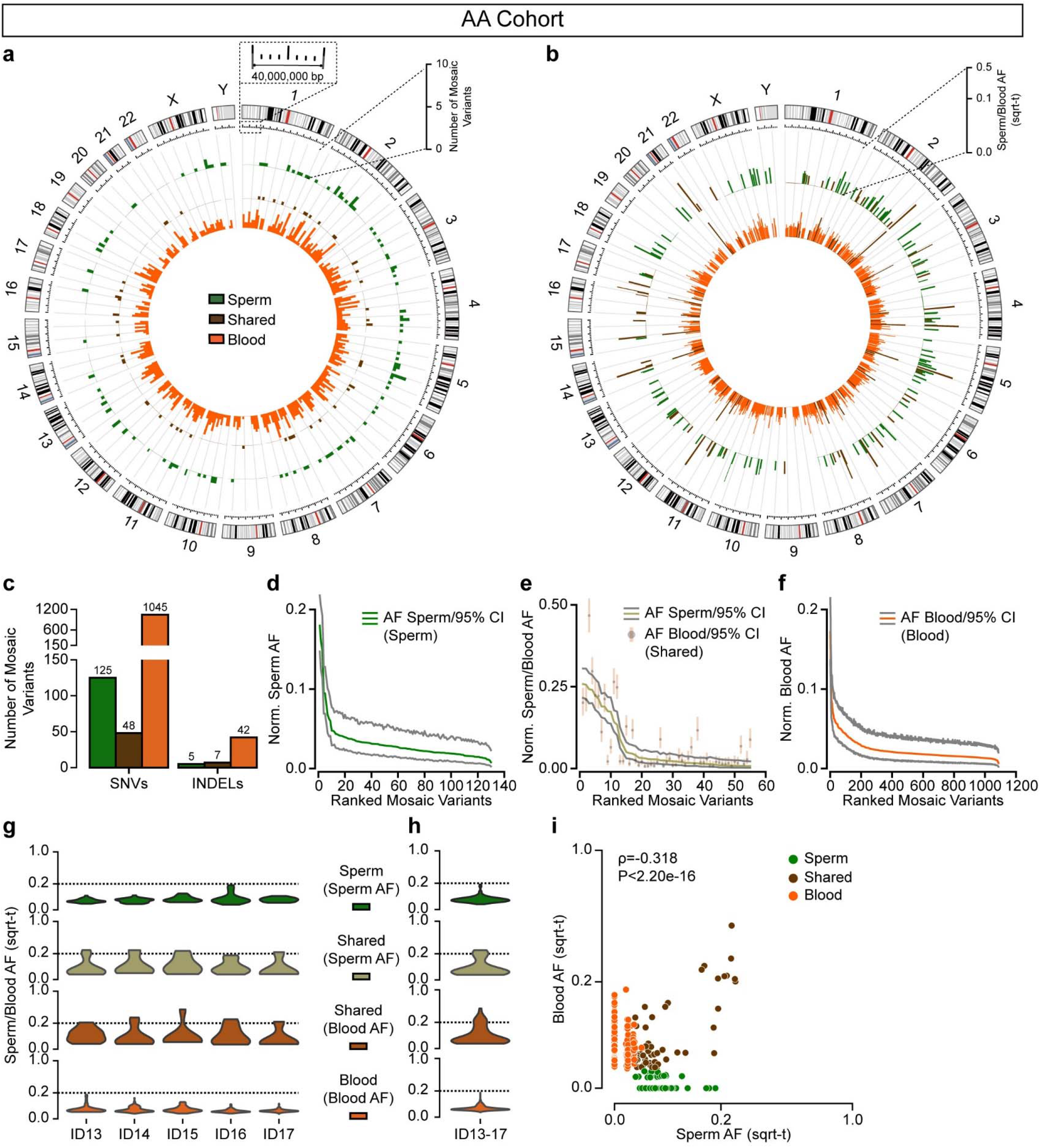
Genomic landscape and AF distributions of mSNV/INDELs detected in the AA cohort. **a**, Circos histograms showing the number of mSNV/INDELs detected from the AA cohort. Color schemes are the same as Fig. 1b. Circles from inner to outer show *Blood, Shared*, and *Sperm* variants. **b**, Circos bar plots of the AFs of mSNV/INDELs and their allelic fractions (sqrt-t on radial axis); the inner circle shows the AFs detected in the WGS of AA blood and the outer circle showed the AFs detected in the WGS of AA sperm. **c**, Number of mosaic SNVs and INDELs detected from the AA cohort. Compared with the YA cohort the number of *Blood* mSNV/INDELs was elevated, in line with the clonal hematopoiesis model. **d**-**f**, Ranked plot of the estimated sperm and blood AF with 95% confidence intervals (exact binomial CIs) from the AA cohort. Other than the number of *Blood* variants, results replicate insights from the YA cohort. **g-h**, AF distribution of *Sperm, Shared*, and *Blood* variants within each individual (g) and the entire AA cohort (h). Consistent with the YA cohort, *Shared* variants show higher peak and overall AF compared to both *Sperm* and *Blood*. i, Correlation between sqrt-t AFs from WGS in blood and sperm of the AA cohort. A significant correlation is observed, similar to Extended Data Fig. 3f. Panel shows Spearman’s ρ and P-value across all variants.

**Extended Data Figure 6.**
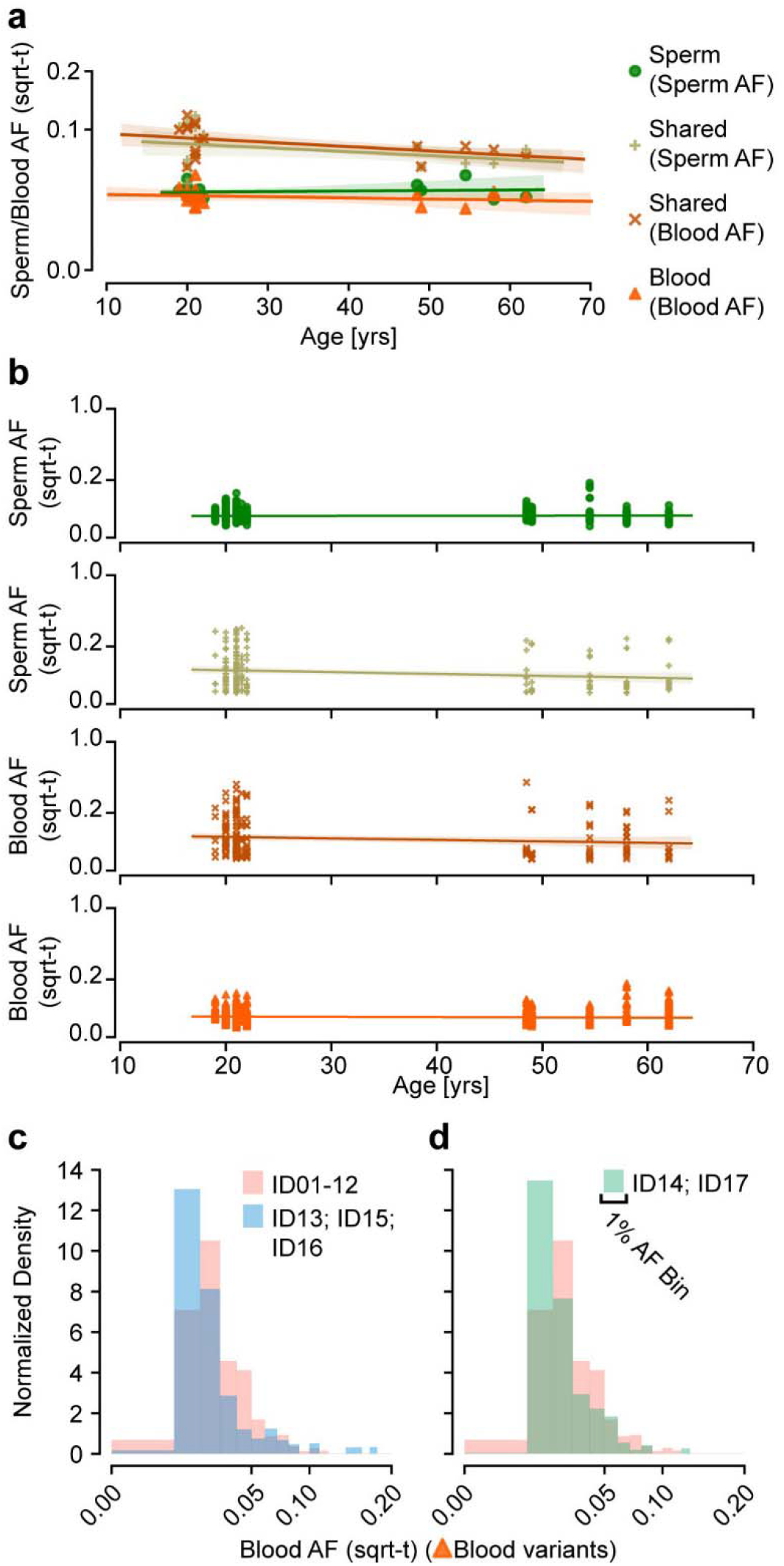
Temporal stability of sperm AFs measured in each individual across YA and AA groups, and age-related clonal- and preclonal-differences between YA and AA blood. **a**, Combined analysis of the average AF of YA and AA mosaic variants relative to the age of individuals. **b**, Each data point represents one variant. y-axis: sqrt-t AFs; x-axis: age at (initial) sample collection. Variants are grouped based on the tissues they were detected from the 300× WGS: AFs of Sperm variants detected in sperm samples, AFs of Shared variants detected in sperm samples, AFs of Shared variants detected in blood samples, and AFs of Blood variants detected in blood samples. Regression lines with 95% prediction intervals are shown for each panel. Results show that the AFs of mosaic variants detected in this study were stable within and between cohorts across a time span of several decades. **c** and **d**, Histogram of the AF distribution of individuals without significant blood clonal collapse (e; ID13, ID15, and ID16) and with clonal collapse (f; ID14 and ID17) compared to YA (ID01-12) individuals. Both sub-groups of the AA cohort exhibited similar differences to the YA cohort despite the difference in *Blood* variant numbers. Panels b and c show the data points, a regression line, and its 95% CI.

**Extended Data Figure 7.**
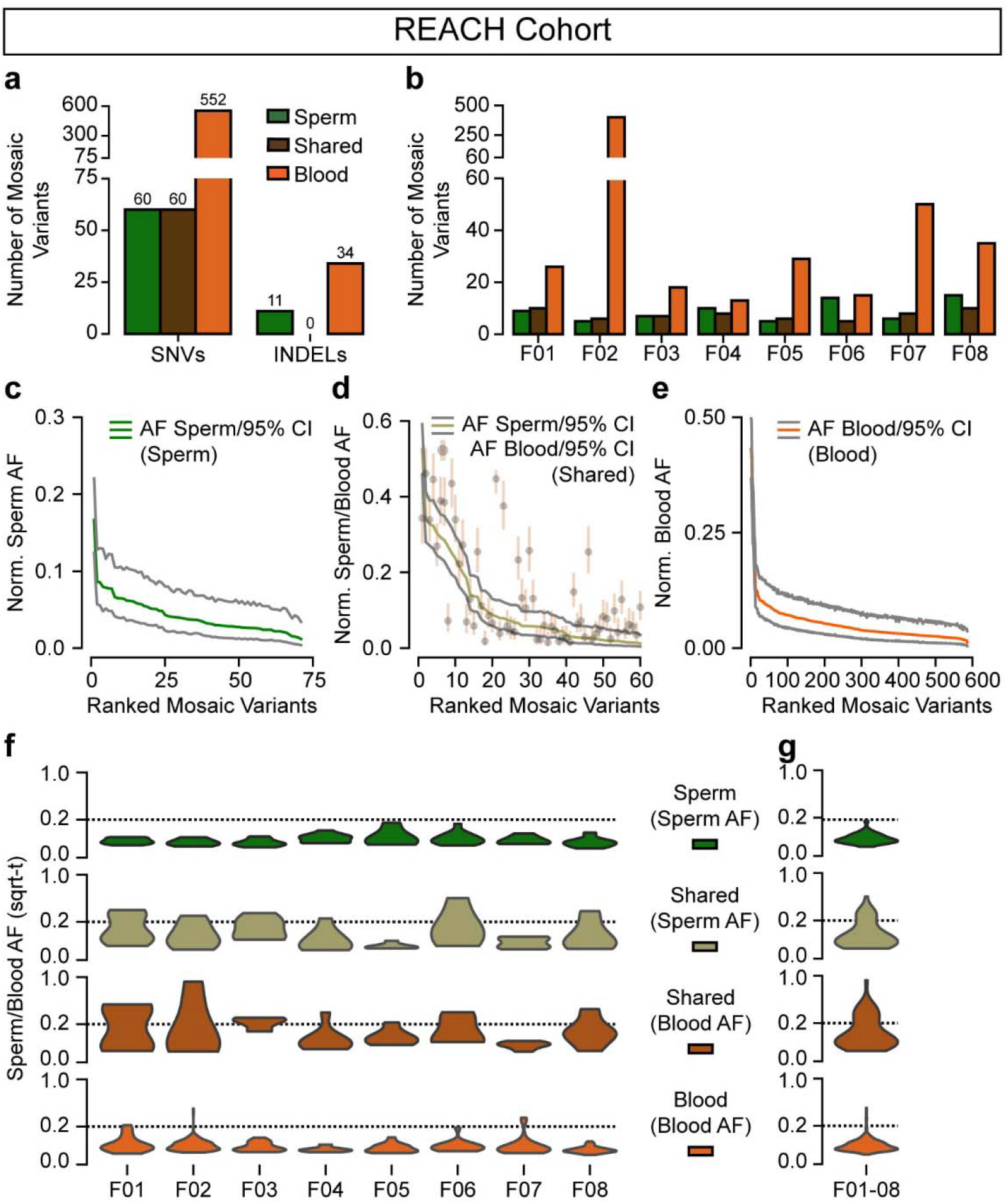
Reanalysis of deep WGS data of sperm and blood from the REACH cohort. **a-b**, Combined (a) and per-individual (b) number of mosaic variants detected in the 8 individuals from the REACH cohort. Compared with the YA and AA cohort the number of mSNV/INDELs detected from the REACH cohort is lower due to the lower sequencing depth. Clonal hematopoiesis was observed in one individual with advanced age (F02; 70 years old). **c-e**, Ranked plot of the estimated sperm and blood AF with 95% confidence intervals (exact binomial CIs) from the REACH cohort. Data reveals similar patterns as for the YA and AA cohorts. **f**-**g**, AF distribution of *Sperm, Shared*, and *Blood* variants within each individual (f) and the entire REACH cohort (g). Consistent with the YA and AA cohorts, *Shared* variants show higher peak and overall AF compared to both *Sperm* and *Blood*.

**Extended Data Figure 8.**
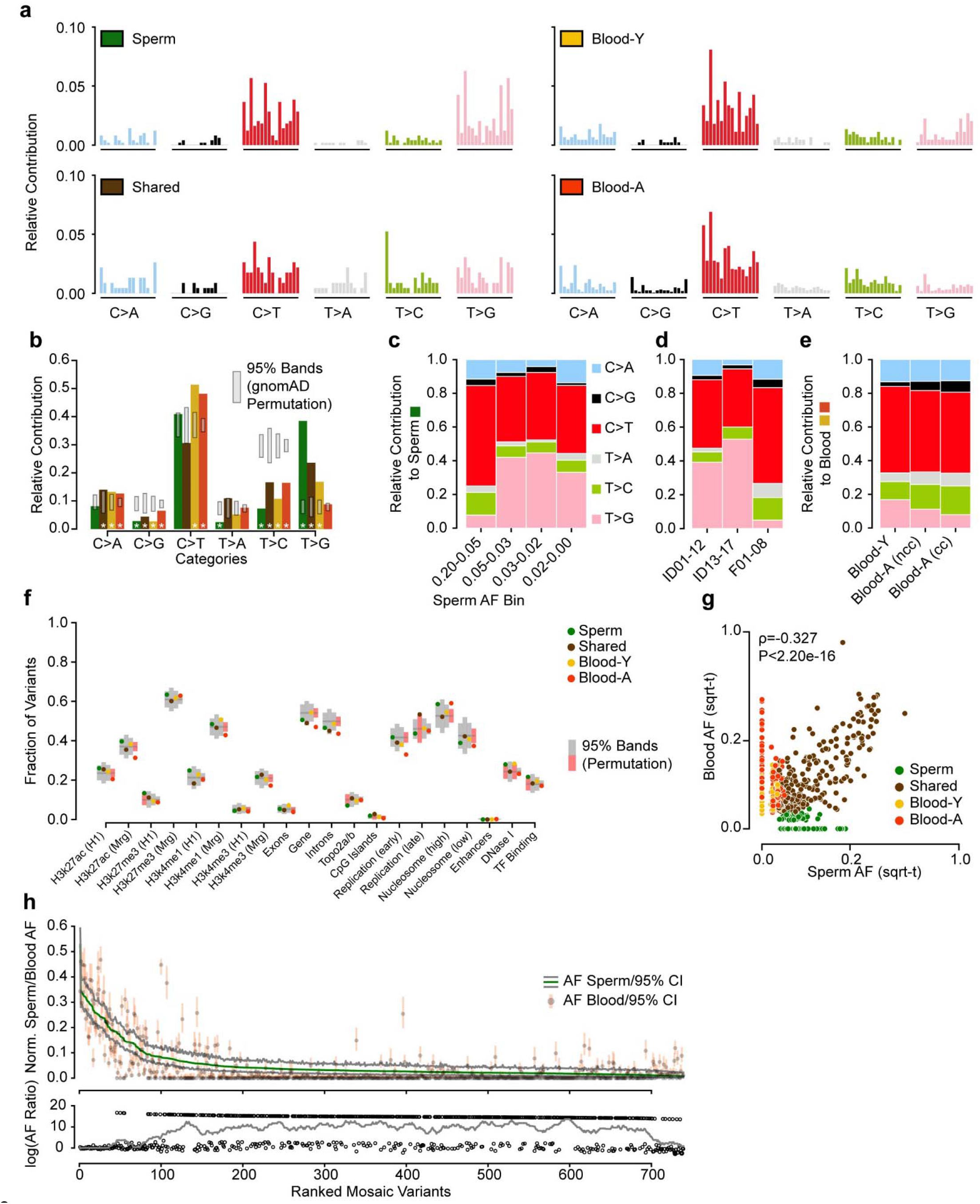
Aggregate analysis highlights base substitution features and distinct mutational signatures. **a**, Base substitution profiles of the 4 different variant groups (tri-nucleotide level). **b**, Signature of the six possible base substitutions of the 4 different variant groups (single nucleotide level). Grey bands: 95% permutation intervals calculated from 10,000 random permutations of gnomAD (v2.0.1) variants using the same number of variants as found from each variant group. Mosaic variants differ from permutations and each other in several categories. Color scheme same as Fig. 4a. **c-e**, Relative contribution of the 6-category base substitution profiles for variants showing C>T predominance and an additional T>G enrichment only in samples with AF lower than 5% in sperm (c), possibly due to the high sequencing depth after break into cohorts (d); YA: ID01-12, AA: ID13-17, REACH: F01-08. C>T relative contribution increased with stronger clonal collapse in blood (e). ncc, non-clonal collapse, cc, clonal collapse. **f**, Fraction of variants located in different genomic regions for the six categories based on tissue distribution. H3k27ac/H3k27me3/H3K4me1 (H1/Mrg): H3k27ac/H3k27me3/H3K4me1 acetylation peak regions measured in human H1esc or merged from 9 different cell lines; Top2a/b: topoisomerase binding regions; Early and Late replication: measured DNA replication timing; Nucleosome (high/low): nucleosome occupancy tendency; Enhancers: annotated enhancer regions; DNase I: DNase I hypersensitive regions; TF Binding: Transcription factor binding sites. 95% permutation intervals were calculated from 10,000 random permutations of the same number of variants from 10,000 random permutations from Simons Simplex Consortium *de novo* variants (if variant residing outside of the permutation interval it is colored red). *Blood-A* showed the most deviations from expectations. **g**, Correlation between sqrt-t AFs from WGS in blood and sperm in each of the four aggregated groups. Panel shows Spearman’s ρ and P-value across all variants. **h**, Ranked plot of estimated sperm and blood AF with 95% confidence intervals for all 773 gonadal mosaic variants detected as mosaic in sperm. Lower plot shows the log_10_ transformed ratio of sperm and blood AFs (0 replaced by 1e-8) and the rolling average over 20 data points to display the local trend. Sperm specific mosaic variants started from an AF of 15% and showed a relatively lower AF.

**Extended Data Figure 9.**
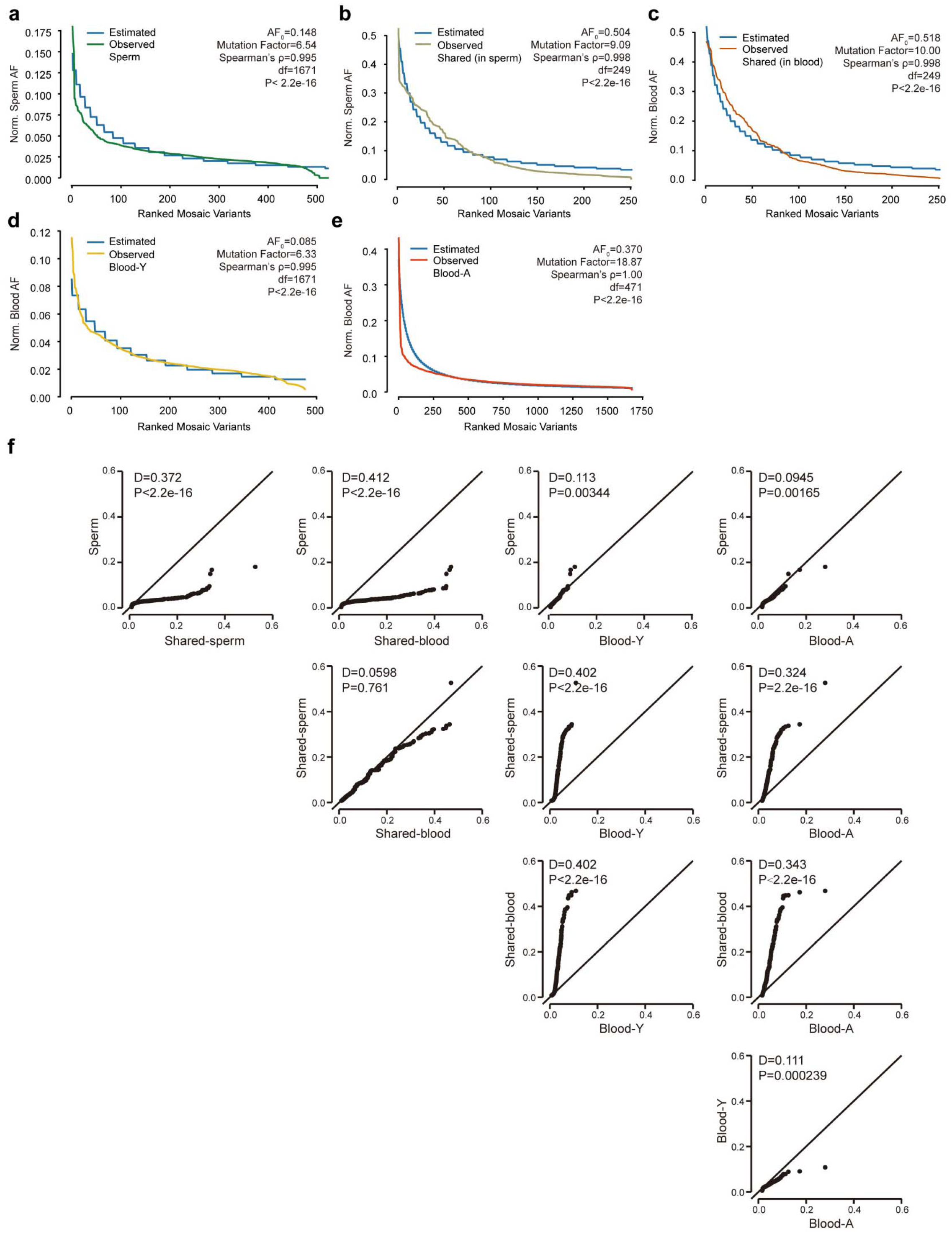
A step-wise exponential regression model to estimate differences in mutational accumulation and quantile analysis of variant classes. **a-e**, Ranked plot of the sex-chromosome normalized AFs measured for variants from each aggregated class; observed values are colored with a color scheme as in Fig. 4a. A step-wise exponential regression model is used to model the distribution of the ranked plot with minimum L1-norm loss; the estimated values are shown in blue. The starting AF (AF0) and the Mutation Factor (MF) defining the relative mutation burden accumulation, as well as Spearman’s correlation coefficients (ρ) and P-values are shown for each variant group. All estimations with Spearman’s ρ≥0.995. Ranked plot of variants in *Sperm* (a) was estimated with a MF=0.153, very similar to MF=0.158 from variants measured in *Blood-Y* (d), indicating similar mutation burden accumulation during tissue development. Ranked plot of variants in *Shared* measured in sperm (b) were estimated with a MF=0.110, similar to the same variants measured in blood (c, MF=0.100), indicating similar but relatively increased (compared to *Sperm* and *Blood-Y*) mutation accumulation speed during early embryonic development. Ranked plot of variants in *Blood-A* were estimated with MF=0.053, different from all other observed variant groups. **f**, Quantile plots (qq-plots) showing the comparison of AF distributions between different variant groups (x and y title). D statistic and P-value from a Kolmogorov-Smirnov test are shown on each comparison pair.

**Extended Data Figure 10.**
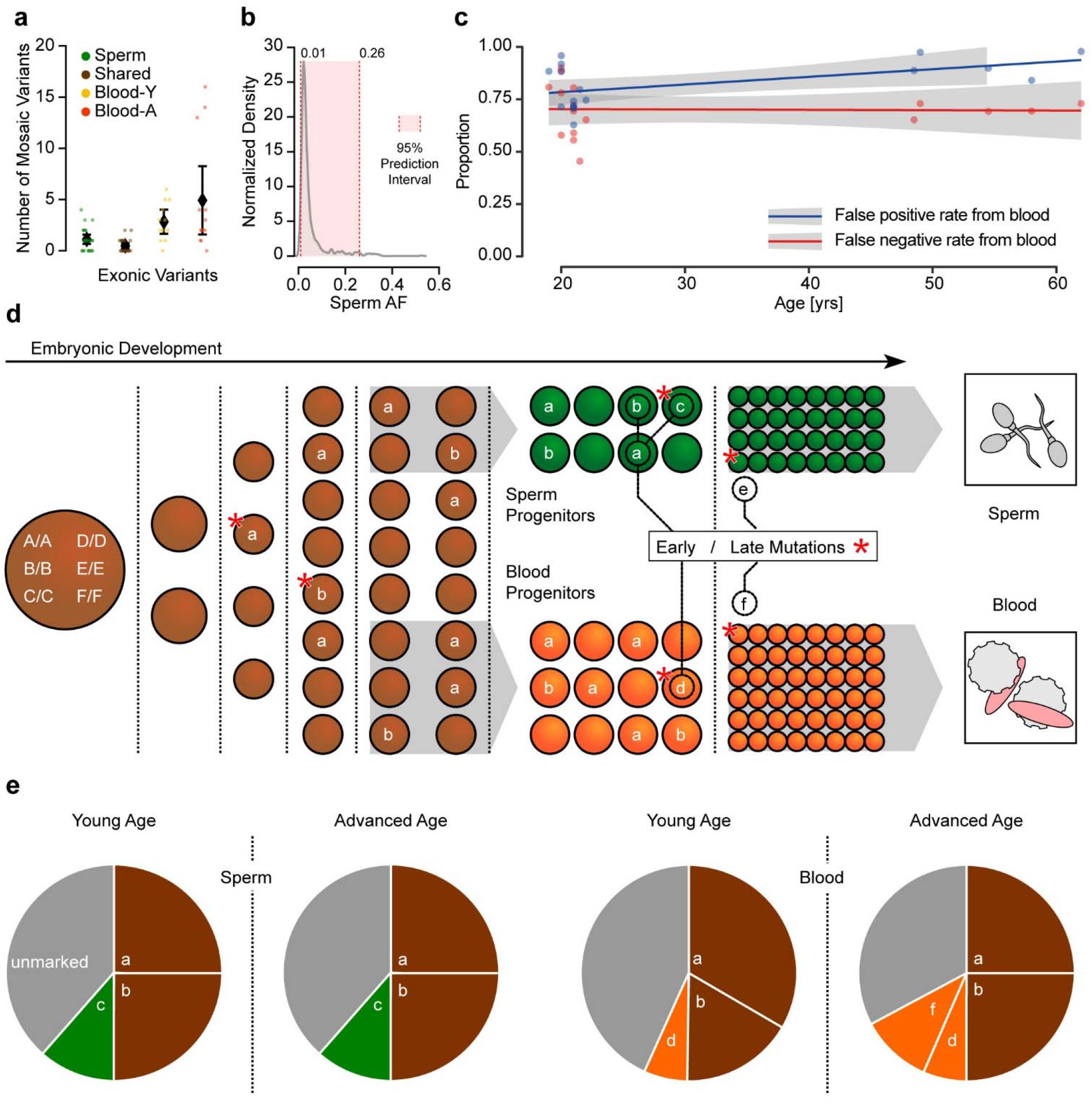
The developmental origin of mosaic mutations and the predictions from the observed temporal variation. **a**, Detectable mosaic variants in each category for exonic variants (c). Shown are individual data points and mean with a 95% confidence interval. **b**, Kernel density estimation of the AF distribution of all sperm mosaic variants. The 95% prediction interval for AF is 1-26%. **c**, Inaccuracy of transmissible mosaicism detection from blood increases with age. Based on the number of blood detectable mosaic variants and their presence in sperm, blood-only detection produces a high false positive rate and the false positive rate is growing with age due to clonal collapse (blue). Blood-only detection produces a consistent 66% false negative rate for the prediction of transmission across different age groups. **d**, Mosaic variants occur throughout development and are typically *Shared* if they occur prior to the split of soma and sperm progenitors. For instance mutation *a* (resulting in genotype *A/a*) occurs during the 4 cell stage, is present in roughly 25% of all cells (i.e. ~12.5% AF), and is shared across blood and sperm. *B/b*, which occurs later, is also shared, but due to stochastic distribution unequally present in sperm and blood. Finally, *C/c, D/d, E/e*, and *F/f* are only present in tissue-specific progenitors, and the latter two happen late and are not detectable in young individuals due to their small AF. **e**, Relative contributions of different variants detected in blood and sperm and their change with age. *Shared* variants are typically present across life, but may change their abundance in blood, due to clonal expansion/collapse. Blood-specific mutations may dynamically increase or decrease with age due to the same phenomenon (e.g. *F/f*). For sperm, this stability across age results in a life-long, predictable transmission risk of variants *A/a*, *B/b*, and *C/c*.

